# Identification of an Essential IL15-STAT1-IRX3 Prosurvival Pathway in T Lymphocytes with Therapeutic Implications

**DOI:** 10.1101/178939

**Authors:** Stephan P. Persengiev

## Abstract

IRX3 homeobox transcription factor plays a role in neural development and in the pathogenesis of obesity and insulin resistance. T lymphocytes play an essential role in the pathogenesis of obesity and their expansion and differentiation is a tightly controlled process. Circulating cytokines trough activation of cell death mechanisms controls the number of CD4 and CD8 T cells, and failure to activate apoptosis is considered the main cause for their uncontrolled expansion. Here, the identification an essential anti-apoptotic pathway in T lymphocytes is presented, whereby IL-15 activation of Stat/Jak signaling cascade leads to induction of the IRX3 expression. IRX3, in turn, promotes T cells survival and self-renewal, at least partially, by controlling the expression of T-bet transcription factor eomesodermin (EOMES). The Stat/Jak signaling inhibition by Jak inhibitor 1 suppresses IRX3 transcription in T lymphocytes and tumor cell lines and increases apoptosis after growth factor deprivation. Our results demonstrate that IRX3 is essential in T lymphocytes differentiation and reveal that the IL-15-dependent IRX3 pro-survival pathway provides potential therapeutic target for regulation of T cell proliferation that has implications for immune cell malignancies, obesity and autoimmune diseases.

## Introduction

IRX3 is a member of the Iroquois homeobox gene family (see IRX1; MIM 606197) and plays a role in an early step of neural development (Bellefroid et al., 1998). In addition, recent findings have implicated IRX3 in the pathogenesis of obesity and insulin resistance (Ragvin et al., 2010; Smemo et al., 2014). In obesity, the white adipose tissue and other organs, e.g. liver and hypothalamus, exhibit a low-grade chronic inflammation (Hotamisligil, 2006; Qatanani and Lazar, 2007), which might contribute to the systemic insulin resistance of type 2 diabetes. Obesity is characterized by the accumulation of diverse immune cell types in adipose tissue (Anderson et al., 2010). The recruitment of macrophages leads to production of pro-inflammatory cytokines that can interfere with insulin signaling (Lumeng et al., 2007). In addition to macrophages, T lymphocytes of the adaptive immune response are recruited to obese adipose tissue. CD8 T cells appear to play a critical role in the development of inflammation and insulin resistance (Jiang et al., 2014b) and the accumulation of CD8+ T cells precede the appearance of pro-inflammatory macrophages (Jiang et al., 2014a; Nishimura et al., 2009). CD8 cytotoxic T lymphocytes are dependent on IL-2 and IL-15 for growth and expansion and recent finding implicated IL-15 in as a negative regulator of visceral fat mass (Carbo et al., 2001). Moreover, IL-15 gain-of-function was associated with lean phenotype, while IL-15 gene inactivation resulted in significant increase in lipid deposition and fat accumulation without alteration in food intake (Barra et al., 2010). These findings and the apparent association of IRX3 with obesity raise the possibility of IRX3 playing a role in the regulatory pathways that govern immune cells responses.

The apoptosis mechanisms that monitor and keep under control the expansion rate of hematopoietic cells are disrupted in a number of human maladies, including autoimmune disorders, obesity conditions, cancer and immunodeficiency (Eguchi, 2001; Hanahan and Weinberg, 2000). Inhibition of apoptosis can be accomplished either by silencing of proapoptotic factors or the activation of prosurvival/antiapoptotic mechanisms that bypass the cell death cascades and has immediate consequences for the immune cell homeostasis (Green, 2003). In the hematopoietic cells, the expansion of activated T lymphocytes is halted by apoptosis activation following the retraction of stimulatory cytokines (Georgopoulos, 1997; Kuo et al., 1997). In several instances, apoptosis induced by growth factor deprivation has been shown to be transcription-dependent (Ishida et al., 1992) and is associated with marked changes in the gene expression profile (Devireddy et al., 2001). Therefore, factors that exhibit differential expression and have a negative effect on cell death/apoptosis programs might be both relevant for normal development and significant for hematopoietic cancers and autoimmunity.

Recently, the upregulation of IRX3 was reported to be linked to the resistance to oncolytic viral therapy in lymphoid malignant cells, in contrast to myeloid malignant cell with low IRX3 expression levels (Lee et al., 2016). Since the activation and deactivation of apoptotic pathways is of particular importance for maintaining immune cell homeostasis and is a primary target of carcingenic insults, we aimed to determine whether IRX3 is involved in this process. We have analyzed the expression of IRX3 in several hematopoietic T cell lineages and determined the effect of interleukin-2 (IL-2) and interleukin-15 (IL-15) deprivation on IRX3 expression. IRX3 expression was found to be dependent on IL-15 and decreased dramatically after IL-15 withdrawal only in CD8 T cells, while remained unaffected by IL-2 removal. These findings identify IRX3 as a factor that plays an essential role in promoting the survival and fate determination of cytotoxic T lymphocytes.

## Results and Discussion

### IRX3 homeobox factor is transcriptionaly activated by IL-15 in T lymphocytes

The gene for the transcription factor Iroquois homeobox 3 (IRX3) that belongs to the TALE/IRO homeobox family and was reported to play a role in neural progenitor patterning (Chen et al., 2011; Rodriguez-Seguel et al., 2009) and have pro-survival functions in human breast cancer (Yang et al., 2010) was repressed by 4-fold after IL-15 withdrawal. To investigate the expression of IRX3 in hematopoietic cells we isolated total RNA from various primary human PBMCs, CD4 and CD8 T cell clones, and several cell lines of hematopoietic and non-hematopoietic lineage and performed semi-quantitative RT-PCR analysis. The results shown in **Figure 1A and B** revealed that IRX3 is expressed exclusively in CD8 T cells, and in actively proliferating transformed hematopoietic and non-hematopoietic lineages. The notable exceptions were Kit-225 cell line that is dependent on IL-2 (Bamford et al., 1994) and CD4 T lymphocytes. In contrast, IRX3 expression was undetectable in primary T cells clones that had limited growth potential and in general proved to be difficult to expand (**Figure 1A**). Thus, we concluded that IRX3 expression is generally associated with the growth factor-dependent mechanisms that support the expansion and self-renewal of T lymphocytes and proliferation of blast-transformed cell lines.

**Figure 1.**
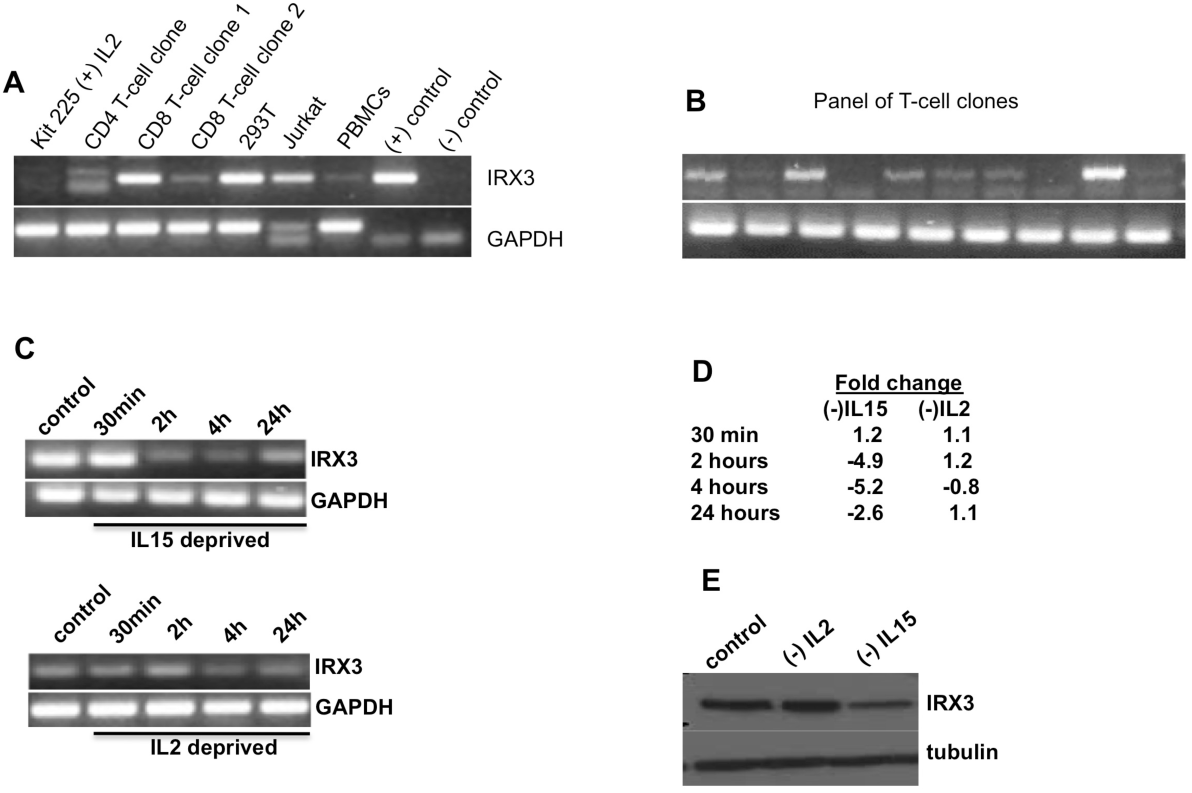
IRX3 is expressed in human T cell lymphocytes and blasttransformed cell lines of hematopoietic and non-hematopoietic origin. **A.** Analysis of IRX3 expression in various cell lines and T cell clones. **B.** Analysis of IRX3 expression in a panel of primary CD8 T cell lines. **C.** Time-course analysis of IRX3 and GAPDH mRNA expression after IL15 and IL2 deprivation. **D.** Quantification of IRX3 mRNA levels following IL2 and IL15 deprivation. **E.** Western blot analysis of IRX3 expression after IL15 deprivation in human CD8 T lymphocytes.

The human cytotoxic T lymphocytes are dependent on IL-2 and IL-15 for growth and expansion. The two cytokines act trough a common membrane receptor and control partially overlapping pathways that are not completely understood (Alves et al., 2007). IL-2 and IL-15 are able to support independently and in combination the survival of primary T cells and IL-2 and IL-15-dependent cells lines. Moreover, each cytokine alone prevents apoptosis activation following either IL-2 or IL-15 withdrawal (**Figure 1**).

Thus, we next explored the cell type specificity of the IRX3 expression and its response to growth factors depletion. Various cell lines and T cell clones were cultured in the presence or absence of the appropriate growth factor (serum or IL-15), and IRX3 expression levels were monitored. RT-PCR analysis revealed that the repression of IRX3 transcripts were first detected in CD8 T lymphocytes within 1 hour of IL-15 withdrawal, but remained unchanged after IL2 deprivation alone (**Figure 1C** and **D**). Similarly, IRX3 protein levels were reduced significantly in CD8 T cells cultured in the abcense of IL15 (**Figure 1E**). Collectively, these data indicate that IRX3 homeobox transcription factor is highly expressed in CD8 T cells and its expression is regulated by the IL15-driven signaling cascade.

### IRX3 acts as a prosurvival factor

CD8 T lymphocytes undergo rapid apoptosis following either IL2 or IL15 deprivation, which raised the possibility of IRX3 to play a role in IL15-mediated CD8 T cells expansion. These results also raised the question whether IRX3 might functions as a prosurvival factor and its downregulation facilitate apoptosis. Thus, in order to verify that IRX3 expression inhibits apoptosis, we transfected CD8 T cells with a N-terminal Flag-tagged IRX3 minigene driven by the constitutive cytomegalovirus (CMV) promoter. IRX3 protein expression was confirmed by the cells with an anti-Flag polyclonal antibody (**Figure 2B**) As expected, the induced expression of an endogenous IRX3 expression vector lowered the apoptotic rate and facilitated cell survival after IL15 deprivation (**Figure 2A**) or treatment with FCC mitochondrial electron electron chain uncoupler (data not shown).

**Figure 2.**
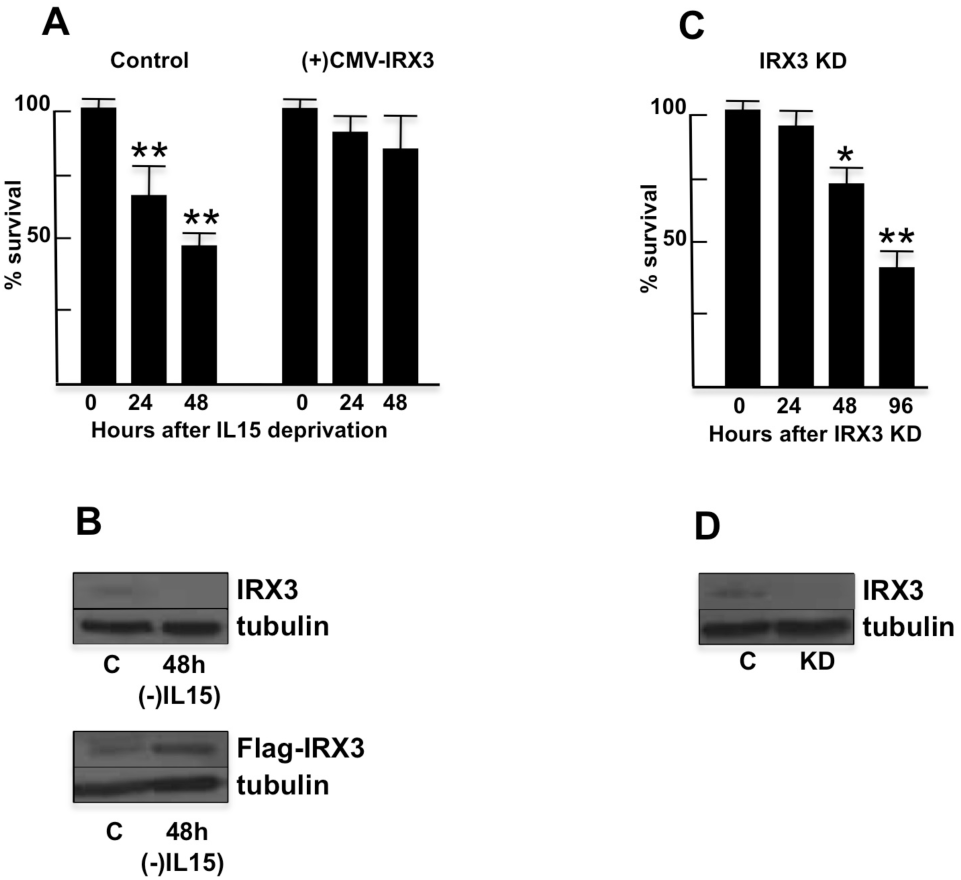
IRX3 induction in derivative CD8 T cells promotes cell survival. **A.** IRX3 overexpression in CD8 T cell inhibits apoptosis. Wild-type and IRX3-T cells were cultured in the presence and absence of IL15 and percentage of apoptotic cells was monitored. **B.** Expression analysis of IRX3 in CD8 T cells expressing CMV-IRX3 minigene. Whole-cell extracts prepared from control and IRX33 cell derivatives were probed sequentially with IRX3 and tubulin antiserums. **C.** IRX3 inhibition by siRNA knockdown reduces the survival rate of CD8 T cells. **D.** Analysis of siRNA-induced IRX3 down-regulation. Whole-cell protein extracts prepared from control and siRNA treated cells were probed with IRX3 and tubulin antibodies. *P<0.05 vs control (C); **P<0.01 vs control (C).

To investigate the possibility that IRX3 downregulation in the cytotoxic T cell clones might lead to activation of apoptosis we knocked down IRX3 by siRNAs in CD8 T cells and monitored the apoptosis response. Loss-of–function experiments in 293T and Jurkat cells performed with IRX3 siRNAs and, as a control, a random non-specific (NS) siRNA, revealed that IRX3 siRNAs substantially reduced the IRX3 levels, whereas the NS siRNA had no effect (**Figure D**). The results shown in **Figure 2C** and **D** revealed that the reduction of IRX3 expression levels by siRNA significantly increased the number of cells undergoing cell death after induction of apoptosis following IRX3 deprivation by IRX3 siRNA transfections.

The results let to the conclusion that IRX3 expression have a significant effect on the rate of cellular proliferation. Taken together, these data indicate that IRX3 expression is essential for the cell survival and is likely to play a role in the mechanisms that support T cell longevity and proliferation capacity.

### IRX3 expression is regulated by Jak-Stat signaling pathway

We next tested whether dependence on IRX3 expression is solely dependent on IL-15 presence or could be modified by other signals that induce apoptosis or act downstream of IL-15 receptors. Inspection of IRX3 promoter region identified response elements for Stat1 and Stat5 transcription factors. Thus, we treated CD8 T cell clones that express IRX3 with either rapamycin or Jak inhibitor 1, which blocked IL-15 signaling trough the TOR and Stat/Jak pathways, respectively, and monitored IRX3 expression. The results of these experiments showed that in primary CD8 T lymphocytes IL15 signaling trough Stat/Jak transcription factors is required to maintain IRX3 expression. (**Figure 3).**

**Figure 3.**
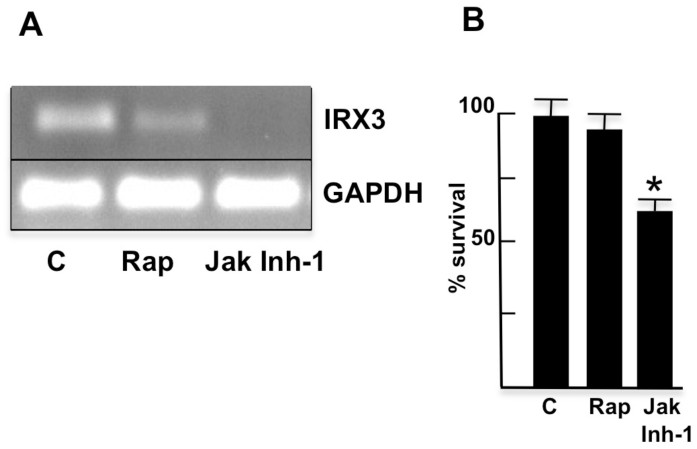
Inhibition of IL15-Stat1 signaling by Jak inhibitor 1 represses IRX3 expression and activate apoptosis. **A.** Effect of rapamycin and Jak inhibitor 1 on IRX3 expression. IRX3 and GAPDH expression was monitored by RT-PCR. **B.** Effect of rapamycin and Jak inhibitor 1 on CD8 T cell survival. *P<0.05 vs control (C).

### CpG islands methylation controls the expression of IRX3

To determine whether reduced IRX3 expression levels in Kit 225 cell line and T cell clones that exhibited accelerated apoptosis rate after IL-2/IL-15 deprivation were the result of epigenetic silencing, we performed bisulfite sequence analysis. **Figure 4B** shows that the IRX3 promoter was densely hypermethylated in Kit 225 cells and CD4 T cell clones, but not in CD8 T cells, Jurkat and 293T cell lines. Treatment of Kit225 cels with the DNA methyltransferase inhibitor 5-aza-2′-deoxycytidine restored IRX3 expression in IRX3 deficient Kit-225 cells, but had no effect in Jurkats and IRX3-expressing T cells (**Figure 4A**). Collectively, these results indicate that loss of IRX3 expression results from epigenetic silencing involving promoter hypermethylation.

**Figure 4.**
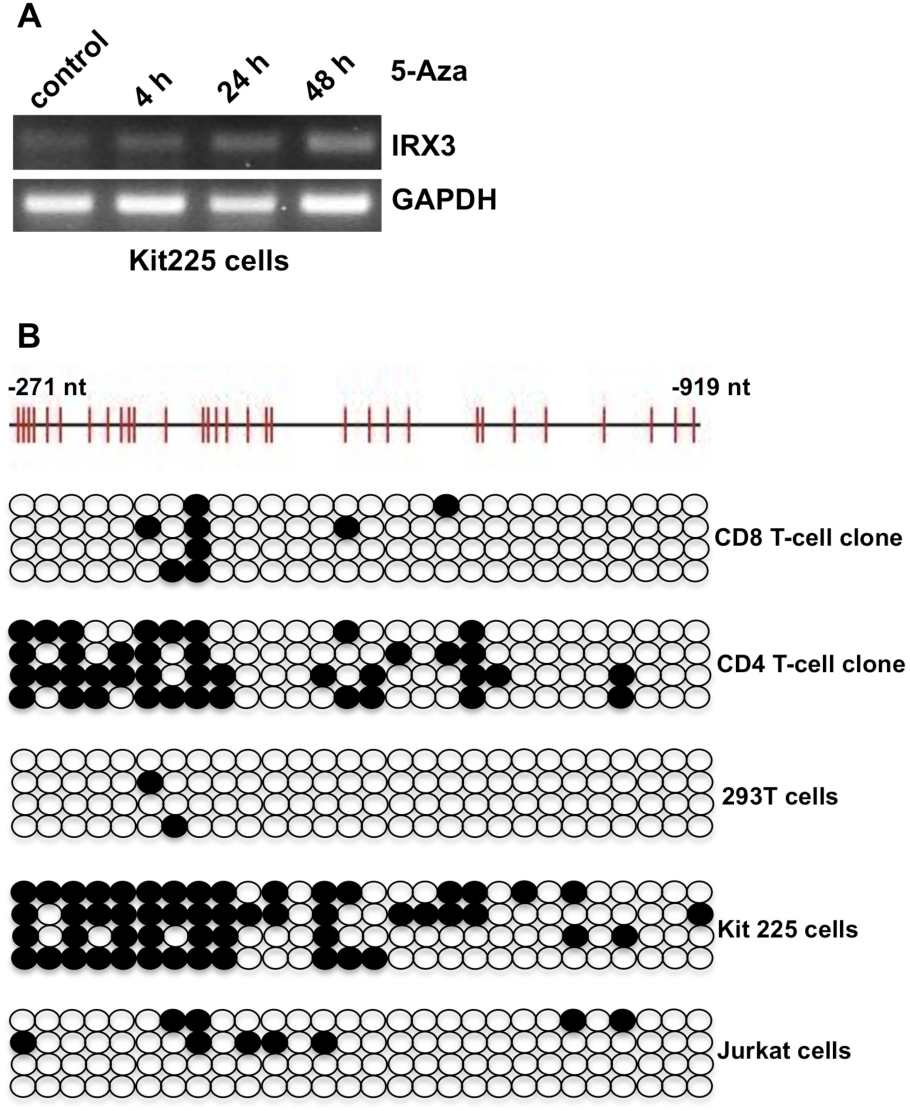
IRX3 promoter is hypomethylated in CD8 T cells and tumorigenic cell lines. **A.** Kit-225 cells were treated with 5-azacytidine (5_Aza) and IRX3 mRNA levels were monitored by RT-PCR. **B.** Bisulfite DNA sequencing analysis of IRX3 promoter in different cell lines and T cell clones.

Importantly, changes in IRX3 promoter methylation after IL15 withdrawal has not been observed indicating that IRX3 downregulation is not the result of epigenetic silencing (data not shown).

### IRX3 regulates EOMES expression

To investigate the in more details the prosurvival activities of IL-15 and identify potential IRX3-regulated transcription nethworks, we isolated total RNA from a primary human pancreatic islet-specific cytotoxic CD8 T cell clones 4h after and IL15 withdrawal and performed a whole-genome transcriptome analysis using Illumina microarray platform. Transcriptional profiles of cells grown in the presence and absence of IL15 were compared. The genes that were regulated selectively by IL15 as compared to the IL15-deprived cells are shown in **Figure 5A**. The expression of OSM, CISH, SOCS1, IRX3, CITED4 and BATF3 genes was selectively reduced in the absence of IL-15, while CEBPD, IL7R7 SQSTM1, EOMES and BCL6 were downregulated. As expected, the gene for the transcription factor Iroquois homeobox 3 (IRX3) was repressed by 4-fold after IL-15 withdrawal. Previous studies have shown that T-bet and eomesodermin (EOMES) transcription factors play a critical role in the proliferative renewal of memory CD8(+) T lymphocytes by inducing the expression of CD122, the receptor that specifies T cell responsiveness to IL-15 (Belz and Kallies, 2010; Intlekofer et al., 2005). This observation raised the possibility that IRX3-mediated T cell survival may be exercised by regulating the expression of BCL6, EOMES or CD122 (IL2R subunit beta). Using transcription factor binding databases (Sandelin et al., 2004), we inspected the promoter regions of BCL6, EOMES and CD122 for potential IRX3 response elements. The CD122 and BCL6 genes was ruled out because they lacked potential homeobox consensus sites, while EOMES contained multiple homeobox response elements within its respective promoters (**Figure 5B)**.

**Figure 5.**
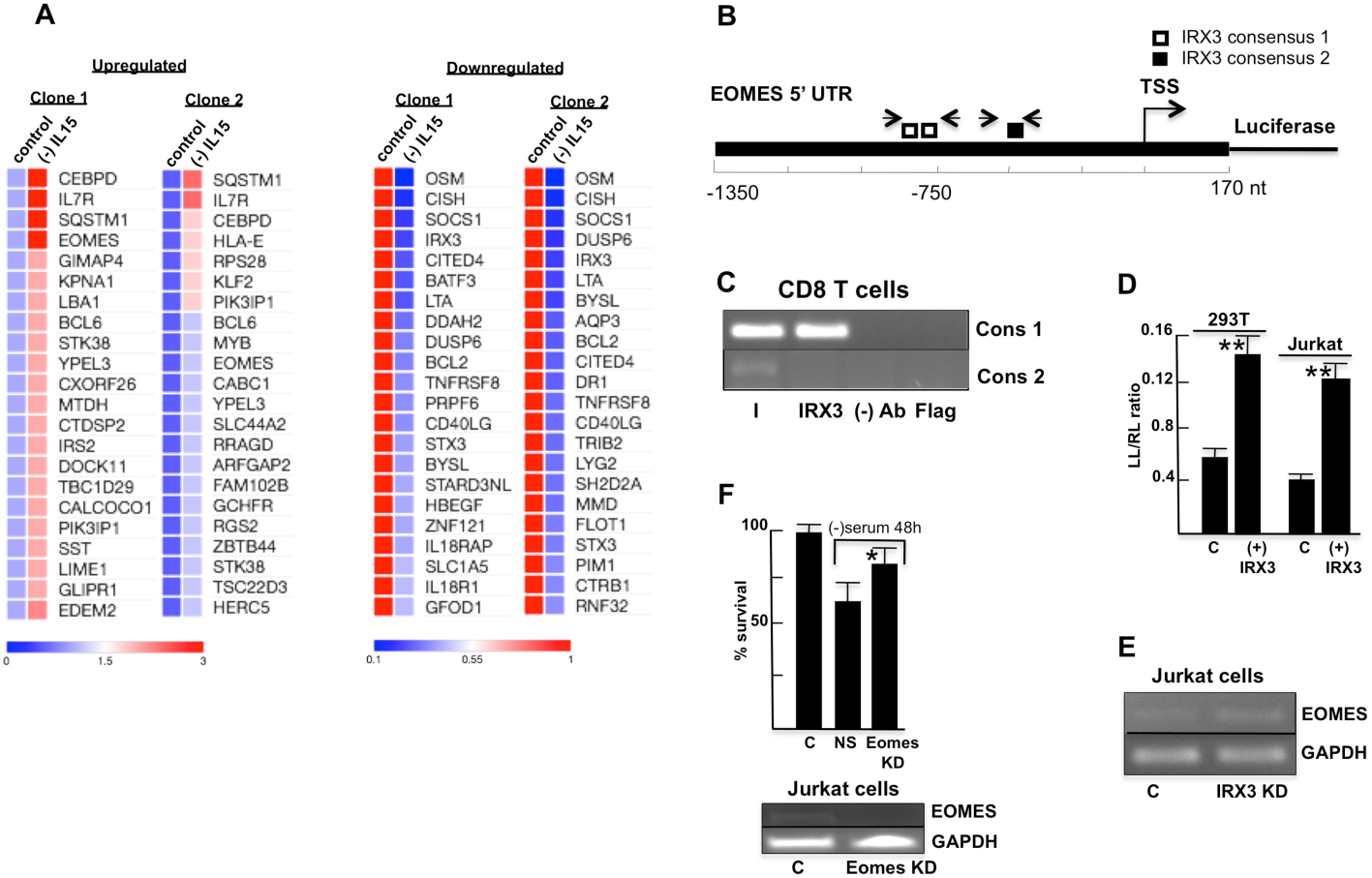
Identification of EOMES as an IRX3 target gene. **A.** Heatmaps of genes with increased or decreased expression in CD8 T cells line after 4 h of IL15 deprivation. Data from two independent experiments are shown. **B.** Schematic of EOMES promoter. Transcription start sites (TSS) and potential IRX3 response elements are indicated. Primer pairs used for chromatin IP analysis are shown. **C.** Chromatin immunoprecipitation (ChIP) analysis of IRX3 association with EOMES promoter in vivo in CD8 T cells. Anti-IRX3 antiserum was used to precipitate the chromatin-bound IRX3, as well as a FLAG antibody was employed a negative control. **D.** EOMES reporter constuct was cotransfected with IRX3 expression vectors and luciferase activity was determined 48 h later in HEK293T and Jurkat cells. **D.** Effect of EOMES knockdown on Jurkat cells survival following serum deprivation for 48 hours. **F.** Effect of IRX3 knockdown on EOMES mRNA expression in Jurkat cells. *P<0.05 vs non specific siRNAs (NS); **P<0.01 vs control (C).

We elected to focus on EOMES for the following reasons: a) EOMES promoter contained two response elements for the homeobox transcription factors NKX2-2, NKX2-5 and NKX6-1 in a favorable sequence context; b) NKX2-2 and NKX6-1 transcription factors have been reported to act in concert with IRX3 in the nervous system, suggesting that the proteins might form functional dimers and share binding sites (Briscoe et al., 2000); and c) EOMES expression was regulated by IL-15. We therefore hypothesized that IL-15, Stat1 and IRX3 comprise a signaling pathway that culminates in regulating EOMES expression. To determine whether IRX3 directly binds EOMES promoter, we preformed ChIP in vivo binding studies using two pairs of primers pairs that spanned potential IRX3 binding sites within EOMES promoter. Increased accumulation of IRX3 was detected at one region of EOMES promoter corresponding to -786 to -740 bp upstream of transcription start site (**Figure 5C**).

Next, we first investigated whether IRX3 was required for EOMES expression. IL-15 deprivation and IRX3 knockdown increased EOMES mRNA levels (**Figure 5A** and **E**). To determine whether the IL15-Stat1-IRX3 signaling pathway promotes EOMES expression at the transcriptional level, we derived a reporter construct in which 1500 bp of the human EOMES promoter was fused to the firefly luciferase gene. **Figure 5D** shows that this EOMES-directed reporter was transcriptionally active when introduced into Jurkat and 293T cells by transient transfections. Interestingly, cotransfection of Jurkat and 293T cells with the EOMES-luciferase reporter and a expression vector containing the entire IRX3 protein-coding sequence under the control of cytomegalovirus promoter (CMV-flag-IRX3) substantially increased EOMES promoter-directed luciferase activity (**Figure 5D).** However, IRX3 knockdown in Jurkat cells led to a notable increase of EOMES mRNA levels (**Figure 5E**). The results demonstrated that EOMES transcription is induced by IRX3, and thus IRX3 can act as a weak transcriptional activator, but is not a direct repressor of EOMES expression in hematopoietic non-hematopoietic cell lineages. The results suggest that the release of IRX3-exerted control of EOMES expression following IL15 deprivation allow the recruitment of a stronger transcription activator to the EOMES promoter, which would account for elevated EOMES transcript levels.

Also, we sought to determine whether EOMES expression is required for cell survival by monitoring Jurkat cells for apoptosis after siRNA-mediated knockdown of EOMES. EOMES knockdown increased substantially the survival rate of Jurkats cells after serum withdrawal (**Figure 5F**). Collectively, these results indicate that EOMES is a *bona fide* downstream effector of IL15-Sta1-IRX3 prosurvival pathway.

IRX3 transcription factor is expressed at high levels in a subset of cytotoxic CD8 T lymphocytes that retain high capacity for proliferation and hematopoietic cells lines derived from T cell tumors. Recent finding gene expression analysis of effector and central memory human CD8 T lymphocytes showed that IRX3 and EOMES are differentially expressed in the different T cell subtypes (Yang et al., 2011). High IRX3 levels were detected in central memory T cells, while EOMES was exclusively expressed in the effector T lymphocytes. These data are consistent with our findings that IRX3 is expressed only in T cell with high proliferative capacity and is regulated by IL-15 and IL-2. Moreover, the data argue that interference with IRX3 function affects the survival and ability of terminally differentiated CD8 T cells to expand after antigen stimulation.

In this study trough a combination of genome-wide gene expression profiling, bioinformatics and reverse genetics, we described a previously unrecognized IL-15-Stat/Jak-IRX3-directed signaling pathway by which EOMES transcription factor is expressed and promotes T cell survival and differentiation. Our results enable us to propose a model for an essential IL-15-regulated anti-apoptotic pathway in human cytotoxic T lymphocytes that promotes T cell proliferation and plays an important role in the development of T cells malignancies. An IL-15-activated Stat/Jak signaling cascade upregulates IRX3. Stat1 directly binds IRX3 promoter, resulting in IRX3 transcription. IRX3 than suppresses EOMES transcription by binding its promoter, resulting in increased cell survival under limited interleukin concentrations (**Figure 6**). It is also worth noting that blast-transformed T cells can produce high levels of IL-2 and IL-15, which in light of the our data would allow them to maintain high proliferation rate by employing a self-sustained autocrine mechanism though constitutive induction of IRX3 transcription (Azzarone et al., 1996; He et al., 2004; Trentin et al., 1996). Our results do not rule out the prospect of additional IRX3 targets, which remain to be identified, to contribute to the inhibition of apoptosis.

**Figure 6.**
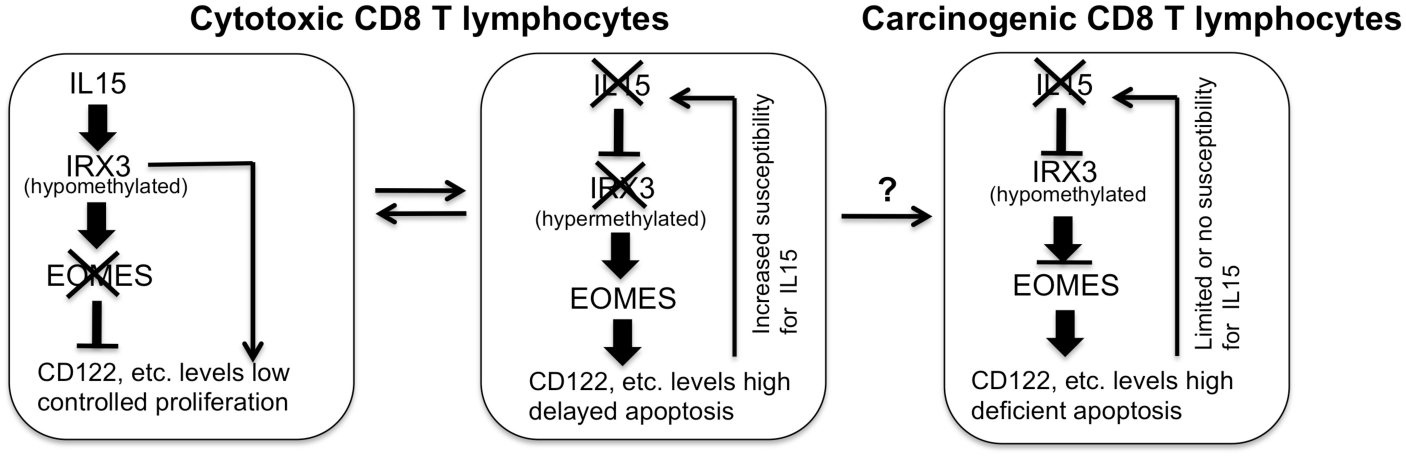
Schematic summary of IL15-Stat1-IRX3-mediated control of CD8 T cell rate of proliferation. IRX3 expression in normal CD8 T cells is controlled b y IL15 pathway, thereby restraining T cell proliferation after IL15 downregulation. In blasttransformed cells IL15-exerted control of IRX3 expression is lost, enabling the cells to escape apoptosis and resulting in uncontrolled proliferation.

These finding suggests that IL15-controled IRX3 transcription pathway is an attractive target for the development of therapeutics for the treatment of obesity-associated conditions, such as type 2 diabetes, and potentially for more effective control of T cells response in autoimmune diseases and T cell cancers. The difficulties associated with development of drugs that target transcription factors (Weiss et al., 2007) in this case can be circumvented by exploring the possibility to target either Jak kinases or IL-2/15 receptors, that are the upstream regulators of IRX3 expression.

## Materials and Methods

### Cell cultures and transfections

The procedures for the isolation of CD3 T lymphocytes and CD4 and CD8 T cell clones from healthy human subjects clones have been previously reported (Bouma et al., 1992). HEK293T, Jurkat and Kit-225 cells blasts were propagated at 37 C in either DMEM or RPMI (Life Technologies) supplemented with 10% fetal bovine serum, 100 U/ml penicillin and 100 U/ml streptomycin. Cells were regularly passed to maintain exponential growth. Human PBMCs, CD3, CD4 and CD8 T lymphocytes were isolated from either type 1 diabetes donors or healthy donors and cultured in Iscoves’s modified Delbucco’s medium (IMDM) supplemented with 10% heat-inactivated human serum (HS), 2 mM glutamine, 100 U/ml of penicillin, and 100 μg/ml of streptomycin (Invitrogen) and IL-2 and IL-15 at 37°C in a humidified atmosphere with 5% CO_2_.

Transfections of HEK293T, Kit225 and Jurkat cell lines with different expression vectors were carried out by using Fugene 6 (Roche Applied Science) according to manufacturer’s instructions. Cells were plated the day before transfection at 2x10^4^ cells per well in 12-well plates. Human T cells transfections were performed by using the Human T Cell Nucleofector Kit according to the manufacturer protocol (Lonza). Luciferase assays were performed 48 hrs later using Dual Luciferase Reporter Assay System (Promega) according to manufacturer protocols. The experiments were carried out in triplicates. IRX3 and EOMES siRNA SMARTpool (Dharmacon) transfections were performed by Oligofectamine (Life Technologies) and 100 nM siRNA mixture, as recommended by the manufacturer.

IRX3 expression vector was constructed by subcloning the full-length human IRX3 cDNA at the C terminus CMV-3xFlag vector. Human EOMES promoter was generated by PCR amplification of 1.7 kb genomic fragment containing EOMES promoter region and subcloned at the N terminus into Luciferase reporter vector. The primers used for IRX3 and EOMES isolations are listed in **Table 1**.

### Cell proliferation assays

The rate of cell proliferation was monitored by using life/dead cell viability **a**ssay(ThermoFisher Scientific) and the trypan-blue live/death exclusion assay protocol.

### mRNA microarray analysis

Total RNA, including miRNA, was purified from cytotoxic T cell clone derived from type 1 diabetes patient and healthy donor by the miRNeasy isolation kit (Qiagen). Total RNA quantification was performed using a NanoDrop N-100 spectrophotometer (NanoDrop, USA) Microarray-based whole genome mRNA expression profiling analysis were performed using Illumina human mRNA arrays (Illumina, USA). Total RNA labeling and hybridization was performed using standard conditions according to manufacturer instructions.

### RT-PCRs

Total RNA from primary CD8 T-cell clones derived from T1D patients and healthy donors was prepared by miRNAEasy kit (Qiagen). cDNA synthesis was carried out with either Superscript II (Invitrogen) and Oligo-dT primers or with TaqMan Reverse transcription kit (Applied Biosystems) according to manufacturer instructions. The sequence of the primers used for detection of human IRX3, EOMES and GAPDH mRNA levels are listed in **Table 1.**

**Table 1.**
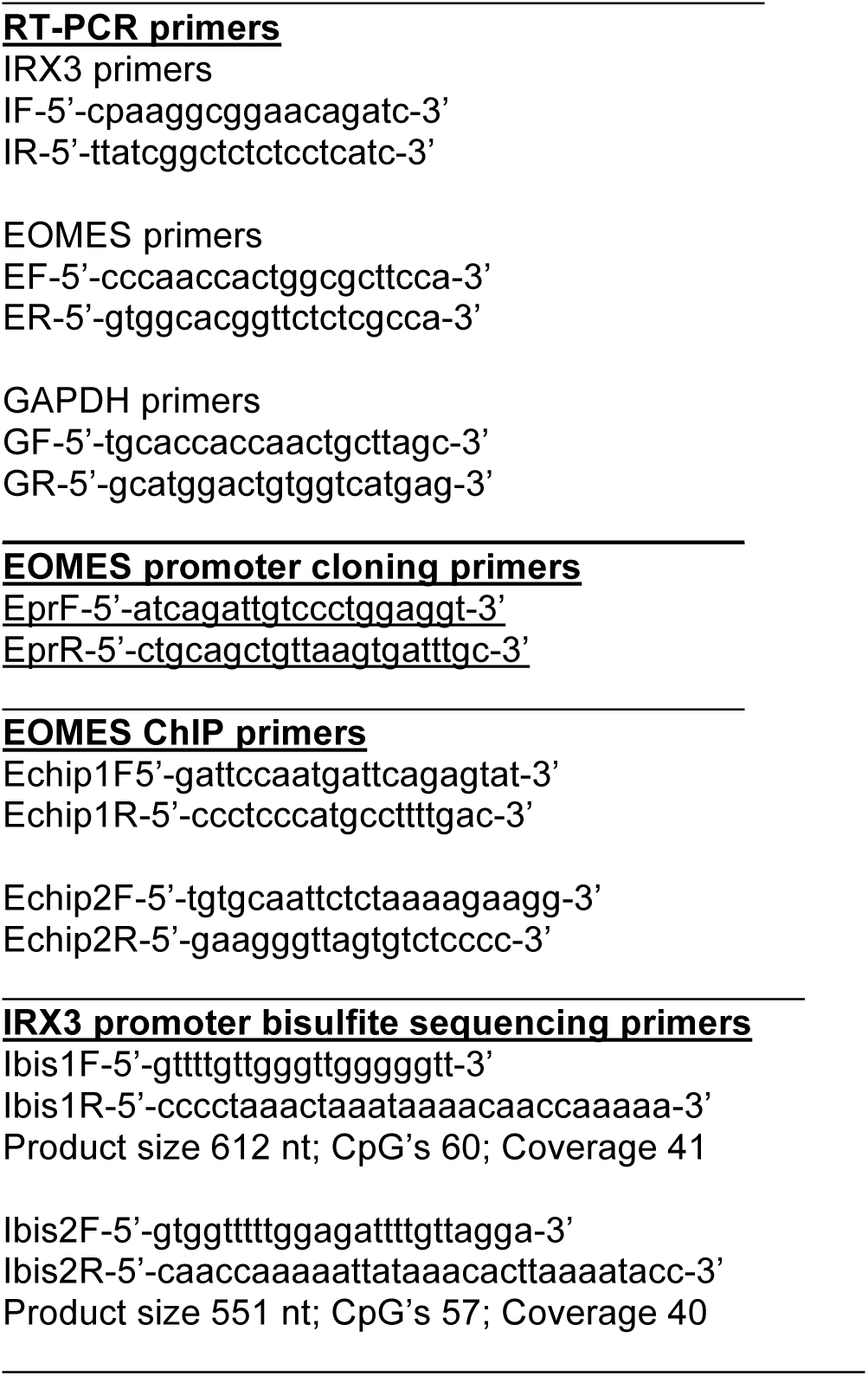
List of primer sequences.

### Protein assay

Total protein isolation was carried out by cell lysis with RIPA buffer supplemented with complete protease inhibitor cocktail (Roche, Branchburg, NJ). Western blot analysis for IRX3 and tubulin were performed with IRX3 and tubulin primary anibodies (Santa Cruz Biotech, Santa Cruz, CA).

### Chromatin immunoprecipitation (ChIP) assays

The amount of IRX3 bound to the endogenous EOMES promoter in CD8 T cells was measured by ChIP assay, using aIRX3 antibody as previously described (Weinmann and Farnham, 2002). Primer sequences used for the PCR amplification of IRX3 promoter fragments containing IRX3 binding site are shown in Table 1.

### Bisulfite sequencing

Bisulfite modification was carried out essentially as described (Frommer et al., 1992). Four clones were sequenced for each cell line (see **Table 1** for primer sequences). For 5-aza-20-deoxycytidine (5-aza) treatment, Kit225 cell lines were treated with 10 mM 5-aza (Calbiochem) for the indicated time points.

### Statistics

Student’s T tests for unpaired samples were performed to compare mean changes in mRNA levels and induced cell death levels.

## Acknowledgements

The experimental results reported in this manuscript were designed by Dr. Stephan Persengiev and performed in the Department of Medical Genetics, UMC Utrecht, The Netherlands. Correspondence

